# Subpopulations in clinical samples of *M. tuberculosis* can give rise to rifampicin resistance and shed light on how resistance is acquired

**DOI:** 10.1101/2025.04.09.647945

**Authors:** Viktoria M Brunner, Philip W Fowler

## Abstract

**Objectives:** Whole genome sequencing (WGS) has become a key tool for diagnosing *Mycobacterium tuberculosis* (M. tuberculosis) infections, but discrepancies between genotypic and phenotypic drug susceptibility testing can hinder effective treatment and surveillance. This study investigates the impact of resistant subpopulations and compensatory mutations in WGS-based rifampicin resistance prediction.

**Methods:** Based on a dataset of 35,538 clinical *M. tuberculosis* samples the sensitivity and specificity of resistance classification were evaluated with and without considering subpopulations and compensatory mutations.

**Results:** By lowering the fraction of reads required to identify a resistance-associated variant in a sample from 0.90 to 0.05, the sensitivity increased significantly from 94.3% to 96.4% without a significant impact on specificity. Allowing compensatory mutations to predict resistance further lowered the false negative rate. Finally, we found that samples with resistant subpopulations were less likely to be compensated than homogeneous resistant samples. Further analysis of these samples revealed distinct clusters with differing amounts of within-sample diversity, pointing towards different mechanisms of resistance acquisition, such as within-host evolution and secondary infections.

**Conclusions:** Our results indicate that a substantial fraction of false negative calls in WGS-based rifampicin resistance prediction can be explained by masked resistant subpopulations. The genetic diversity within the heterogenous samples is consistent with at least 28% of the rifampicin resistance arising from secondary infections.

## Introduction

Tuberculosis is estimated to be responsible for 1.25 million deaths in 2023 with 10.8 million people developing disease ^1^. Resistance to antibiotics is particularly concerning with multi-drug resistant tuberculosis (MDR-TB) estimated to cause 150 000 deaths that year despite only 400 000 people developing MDR-TB ^1^. This is why the first step towards successfully treating *M. tuberculosis* infections is fast and reliable diagnostics, including drug susceptibility testing (DST). Phenotypic DST involves culturing the bacteria in the presence of different antibiotics ^2^ and is reliable for most drugs. However, since *M. tuberculosis* grows slowly, it is time-consuming despite efforts to reduce culture time. ^3^ Genotypic DST instead uses the presence of genetic variants to infer whether a sample is resistant and can be faster than phenotypic DST. ^4^ The prerequisites are that the underlying mechanism is genetic and the exact genetic variants conferring resistance are known. If true, one can then look for the presence of specific genetic variation using rapid molecular tests such as GeneXpert MTB/RIF ^5^, or use whole genome sequencing (WGS) to scan the entire bacterial genome for resistance-associated variants (RAVs) for different antibiotics. ^6^ The latter requires WGS of the patient-derived sample, followed by application of a high-confidence catalogue of RAVs, enabling the sample to be classified as susceptible or resistant to a panel of antibiotics. ^7^ Due to its cost and complexity, WGS will not become a front-line diagnostic method in many high-burden countries in the near-future but could fill the need for ongoing national surveillance to identify resistant strains in circulation not detected by cheaper molecular tests. ^8^

The second edition of the WHO catalogue of mutations contains the most comprehensive list of RAVs to date for predicting rifampicin resistance in *M. tuberculosis* and achieves 93.3% sensitivity and 96.9% specificity on its training dataset compared to standard phenotypic DST results. ^9^ Despite being one of the best-performing drugs, the sensitivity for rifampicin resistance prediction remains below the 95% threshold proposed for antimicrobial susceptibility test devices by the International Standards Organization (ISO). ^10^ It is, of course, unrealistic to expect perfect agreement between the results of phenotypic and genotypic DST but any discrepancy creates problems not only for scientific efforts investigating disease transmission networks of *M. tuberculosis*, ^11^ but also for diagnostics. ^12^ If WGS-based DST is to complement or even replace phenotypic DST in some settings, it needs to achieve comparable performance. Hence there is a strong need to close the gap between phenotypic and genotypic DST results.

Here, we show the performance of WGS-based DST for rifampicin can be trivially improved by permitting resistant subpopulations to contribute to the final classification. This has already been shown to significantly improve the sensitivity of resistance prediction for fluoroquinolones in *M. tuberculosis*. ^13^ The hypothesis is that the genotypic approach is under-calling resistance since resistant subpopulations are not identified by many WGS bioinformatics pipelines, yet phenotypic DST methods will flag samples as resistant if as little as 1% of the bacteria are resistant ^14^. The phenomenon of heteroresistance has been described in many pathogens. ^15^ However, for most pathogens, resistant subpopulations are not expected since the standard laboratory process is to pick single colonies, constraining the detection of any heteroresistance. This is not true for *M. tuberculosis* where e.g. an aliquot taken for DNA extraction from a BD Mycobacteria Growth Indicator Tube (MGIT™) tube usually contains multiple ‘crumbs’ ^16^ and so could readily harbour resistant subpopulations, reflecting more accurately the patient’s infection.

We define the fraction of read support (FRS) as the proportion of reads at a genetic locus that supports a specific genetic variant. Bioinformatics pipelines conventionally specify a minimum FRS for a genetic variant to be called. The first edition of the WHO *M. tuberculosis* mutation catalogue ^17^ and the CRyPTIC consortium ^18^ both used a conservative FRS threshold of 0.90 when calling genetic variants. This high threshold prevents sequencing errors leading to spurious variant calls, but as sequencing technologies have improved and error rates decreased, this now seems unduly conservative since it also ensures genetic subpopulations are not detected. Such subpopulations are expected if an infection has only recently evolved resistance, perhaps in response to treatment, or if there has been a secondary resistant infection.

A second way to improve the performance of catalogue-based predictions is to include more resistance-associated variants. For now, catalogues only contain alleles that (are assumed to) directly cause resistance. We show that including genetic variants indirectly linked to resistance, such as compensatory mutations (CMs), can improve sensitivity without lowering specificity. CMs arise in response to a reduction in fitness caused by rifampicin resistance mutations and hence whilst not causal are only found in rifampicin-resistant samples. ^19–21^ Hence including them in a catalogue would mainly be useful for samples where the RAV cannot be resolved, such as those with low sequencing quality. We have previously identified a list of high-confidence CMs using their strong association with the presence of resistance mutations. ^22^ We aim to test if including this list in the official WHO resistance catalogue ^9^ improves the performance metrics of resistance prediction.

Samples with resistant subpopulations capture an interesting state in *M. tuberculosis* infections which can arise via multiple different scenarios (Fig. 1). The first possibility is within-host evolution of resistance, where the resistant subpopulation gradually takes over a susceptible population during drug treatment (Fig. 1A). Secondly, the evolved drug resistant subpopulation can revert to a susceptible phenotype or be outcompeted after drug treatment is stopped (Fig. 1B); or thirdly the resistant subpopulation is acquired through a secondary infection (Fig. 1C). Distinguishing between these possibilities requires that we know whether the sample was taken before, during or after drug treatment. Unfortunately, precise data on this is scarce in WGS datasets and hence we can seldom distinguish between scenarios 1 and 2 (Fig. 1A, B). The second unknown variable is the source of resistance: within-host evolution or a secondary infection. We aim to discern the source of resistance by quantifying the amount of genetic diversity in the samples with resistant subpopulations in our dataset.

**Figure 1:**
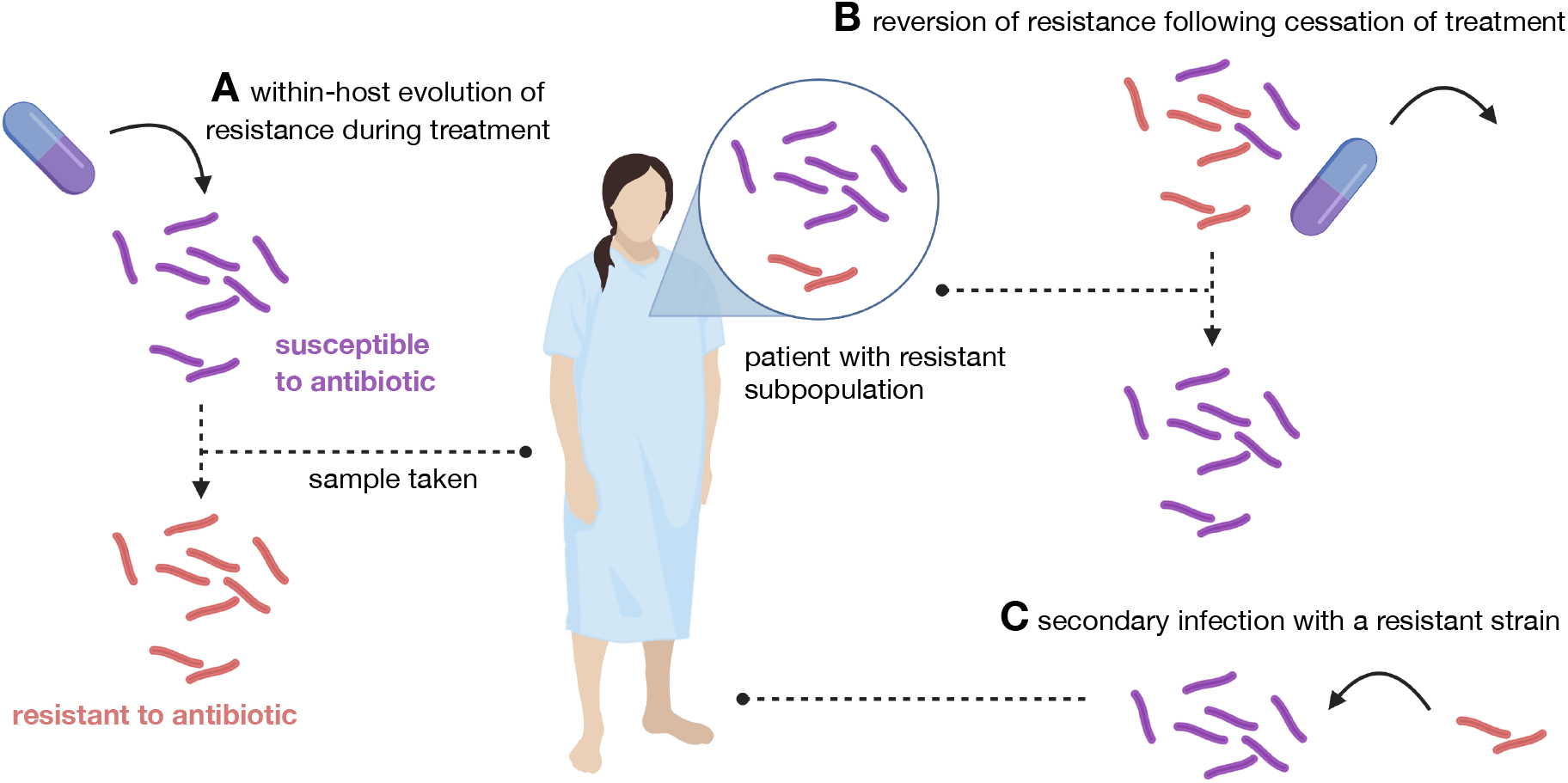
Three ways a patient can acquire an infection with a resistant subpopulation. The susceptible subpopulation is shown in purple, the resistant subpopulation in red. Samples with resistant subpopulations can originate from multiple different scenarios, among them (**A**) within-host-evolution,(**B**) reversion to susceptible phenotype following cessation of treatment, and (**C**) secondary infection.

## Materials and Methods

### Dataset sources

In addition to collecting *>*20,000 *M. tuberculosis* samples, each of which underwent whole genome sequencing (WGS) and drug susceptibility testing (DST) using one of two bespoke 96-well broth microdilution plates, ^18;23^ the Comprehensive Resistance Prediction for Tuberculosis: an International Consortium (CRyPTIC) project also aggregated *M. tuberculosis* samples with WGS and/or DST data that had been previously published. An earlier version of this dataset with heterogeneous DST methods was used to build the first edition of the WHO catalogue of tuberculosis resistance-associated variants. ^24^

### WGS data processing

A subset of 41,575 publicly available samples from the European Nucleotide Archive with short-read paired-end FASTQ files (version 3.0.0 of the CRyPTIC dataset ^25^) were uploaded to the EIT Global Pathogen Analysis Service (GPAS, https://gpas.global) and processed using version d5f9cd0 of the Mycobacterial pipeline. ^26^ The variant caller, Clockwork, calculates the ‘fraction of read support’ (FRS) and, by default, only calls genetic variants where FRS *≥* 0.90 with a filter applied to the remainder. ^27^ Here, we instructed the downstream tool, gnomonicus, to ignore these filters and record all potential genetic variants so that we could detect subpopulations of bacteria. ^28^ Most samples (37,594) had one or more of 101,020 mutations detected in the RNA polymerase (genes *rpoA, rpoB, rpoC, rpoZ* and *sigA*). This includes 3,260 so-called null mutations where there are insufficient (*<* 3) reads to call a variant. To protect against spurious calls due to sequencing errors, variants needed to be supported by at least three reads.

Examining *rpoB* in more detail we find 31,347 samples containing 58,789 mutations with a FRS *≥* 0.90 and 1,372 samples containing 1,940 mutations supported by an FRS *<* 0.90. A small number of the latter samples have very large numbers of putative mutations, indicative of contamination or poor sequencing quality. In both cases the most common *rpoB* mutations were A1075A and S450L, as expected. Mutations were flagged as being associated with resistance to rifampicin according to the second edition of the WHO catalogue of mutations in *M. tuberculosis*. ^9^ We used a published list of high-confidence compensatory mutations ^22^ to annotate which mutations are compensatory.

### Phenotypic DST data processing

A total of 52,148 samples had one or more binary rifampicin drug susceptibility test results using a range of methods; the most common were broth microdilution plate (24,172 samples) and mycobacterial growth indicator tube (23,682). If a sample had been tested using more than one method and all methods produced the same S/R result then, if present, the CRyPTIC result was retained since it has richer data and has undergone additional quality control. If the methods disagreed on the outcome then the first resistant result was retained. This resulted in 48,031 samples with a single rifampicin DST result. Merging with the genetic data identified 35,538 samples which have both whole genome sequencing data and a single rifampicin DST binary result: this dataset forms the basis of our subsequent analysis. The sensitivity and specificity of the genotypic resistance call were calculated with respect to the phenotypic drug susceptibility result. Significance was tested using a proportions z-test.

### Reproducibility statement

All analysis, figures and tables in this paper can be reproduced using a GitHub repository which contains all data tables and code. ^29^

## Results

### Classifying subpopulations that contain rifampicin RAVs as resistant improves sensitivity

In our dataset of 35,538 samples, we will describe samples with a rifampicin resistance-associated variant (RAV) with a fraction of read support (FRS) *≥* 0.90 as *homogeneous*, and samples containing RAVs at an FRS *<* 0.90 as *heterogeneous*. Mutations with an FRS *<* 0.90 read support will be described as *minor*. Out of the 10,568 samples containing a rifampicin RAV at any level of read support, 10,287 are homogeneous, 261 are heterogeneous and 20 are mixed, i.e. contain at least two rifampicin RAVs, one supported by FRS *≥* 0.90 and another with FRS *<* 0.90 (Fig. 2A). Examining the distribution of minor rifampicin RAVs, we observe that RAVs are seen down to an FRS of 0.085, possibly increasing at higher values of FRS (Fig. 2B). To detect a minor RAV at an FRS of 0.05 necessitates a read depth of 60 since three reads are required to call a minor RAV to avoid false positives; this is satisfied, on average, for 79.8% of the dataset (Fig. S3). This indicates that the full range of values should be considered when assessing the impact of the FRS threshold on the performance of resistance prediction.

**Figure 2:**
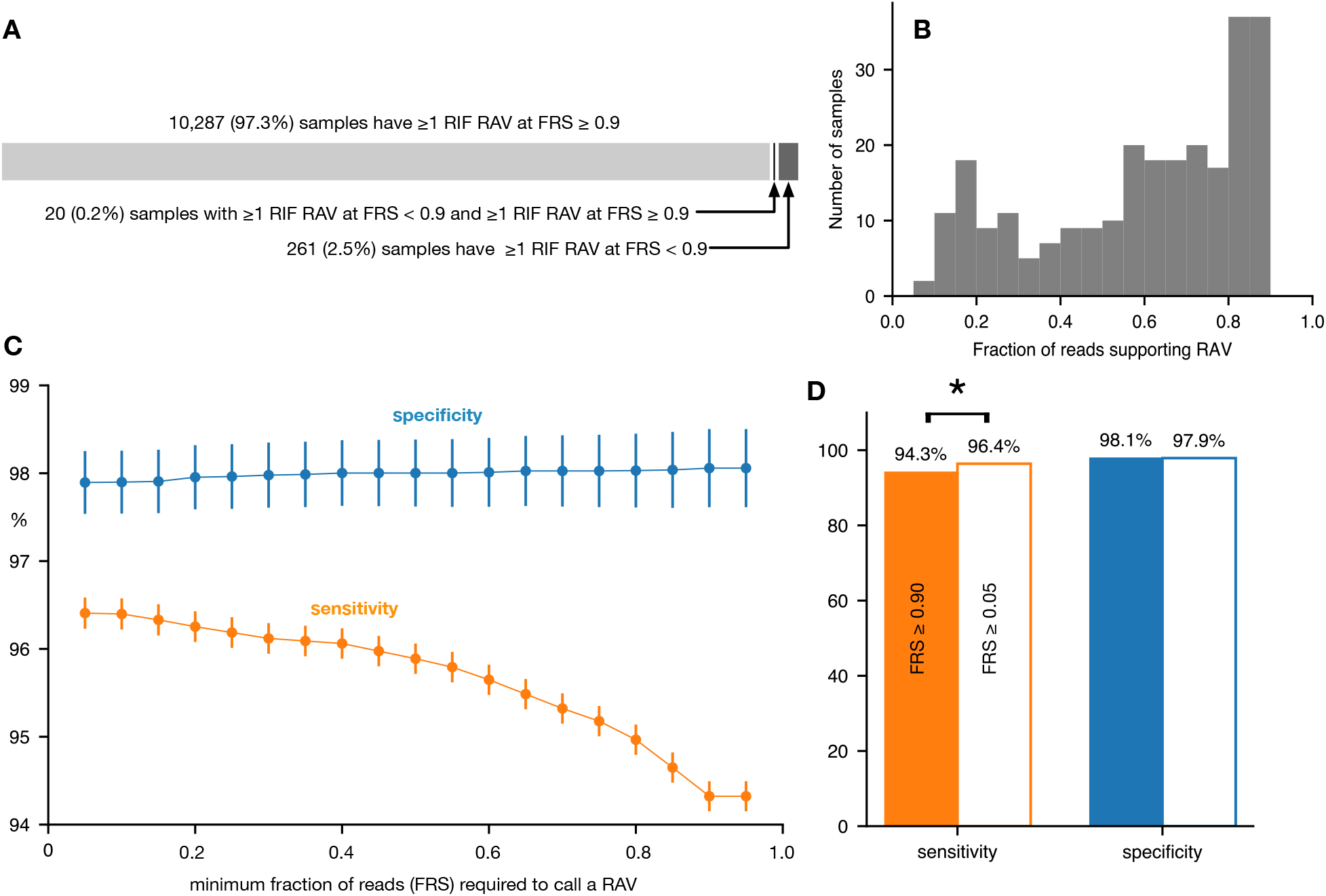
Lowering the fraction of read support (FRS) threshold for calling rifampicin (RIF) resistance-associated variants (RAVs) increases sensitivity with no significant effect on specificity. **(A)** The majority of samples containing a RAV are homogeneous, meaning they show a RIF RAV at *≥* 0.90 FRS. The heterogeneous samples make up for 2.5% of samples, but we rarely see samples with both a RAV at ≥ 0.90 FRS and a RAV with lower read support. **(B)** The distribution of FRS for the RAVs in the 258 heterogeneous samples with 0.05 ≤ FRS *<* 0.90. **(C)** Decreasing the FRS threshold required to support a variant call that is a known RAV increases sensitivity of the prediction with little effect on specificity. The slopes of a linear regression for sensitivity and specificity are -0.02 and 0.002, respectively. Error bars (95% confidence limits) are plotted as calculated via the binomial proportion. **(D)** Sensitivity is significantly improved if the FRS threshold is lowered from 0.90 to 0.05 (z-test, p-value = 8e-13). There is no significant change in specificity. For clarity error bars are not plotted.

The sensitivity increases from 94.3% to 96.4% as the minimum FRS required to call a RAV is decreased from 0.90 to 0.05; over the same range the specificity falls slightly from 98.1% to 97.9% (Fig. 2C, Table 1). The increase in sensitivity when the FRS is dropped from 0.90 to 0.05 is statistically significant, whilst the decrease in specificity is not (Fig. 2D). Over the same range the negative predictive value (NPV) also significantly increases whilst there is no significant change in the positive predictive value (PPV, Fig. S1, Table 1). Taken together, this indicates that applying a conservative FRS threshold masks resistant subpopulations, resulting in falsely predicting some samples as susceptible to rifampicin.

**Table 1:**
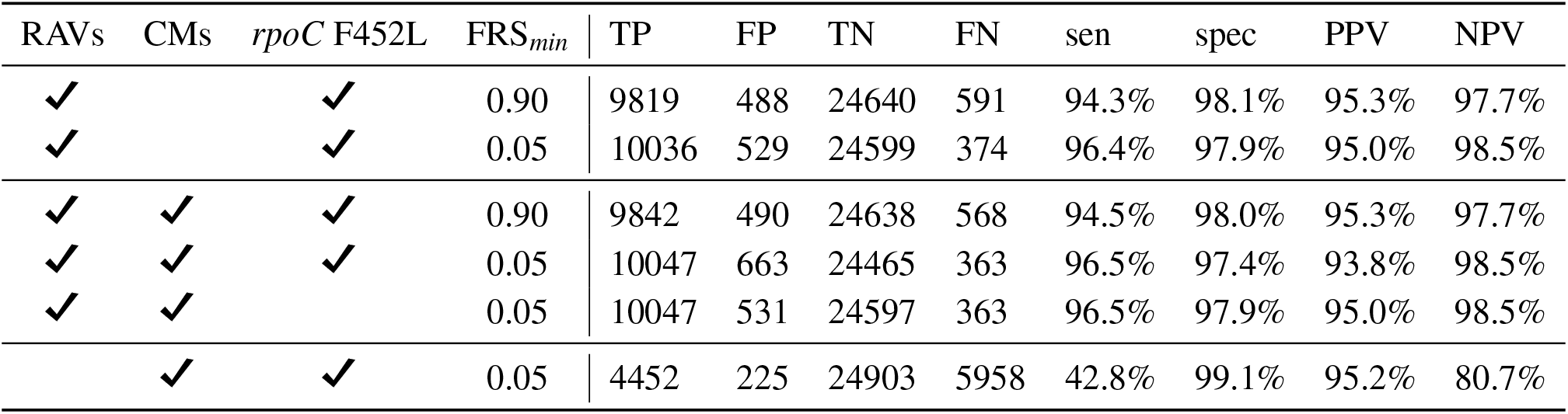
Fraction of read support (FRS) and corresponding contingency table values and performance metrics for different scenarios of the catalogue-based predictions for rifampicin resistance. Scenarios are shown for different FRS thresholds, and are either using RAVs and/or CMs for prediction or not. In one scenario, the CM *rpoC* F452L was removed from the analysis. “sen” shows sensitivity and “spec” the specificity of the resistance prediction.

| RAVs | CMs | *rpoC* F452L | FRS<sub>min</sub> | TP | FP | TN | FN | sen | spec | PPV | NPV |
| --- | --- | --- | --- | --- | --- | --- | --- | --- | --- | --- | --- |
| ✓ |  | ✓ | 0.90 | 9819 | 488 | 24640 | 591 | 94.3% | 98.1% | 95.3% | 97.7% |
| ✓ |  | ✓ | 0.05 | 10036 | 529 | 24599 | 374 | 96.4% | 97.9% | 95.0% | 98.5% |
| ✓ | ✓ | ✓ | 0.90 | 9842 | 490 | 24638 | 568 | 94.5% | 98.0% | 95.3% | 97.7% |
| ✓ | ✓ | ✓ | 0.05 | 10047 | 663 | 24465 | 363 | 96.5% | 97.4% | 93.8% | 98.5% |
| ✓ | ✓ |  | 0.05 | 10047 | 531 | 24597 | 363 | 96.5% | 97.9% | 95.0% | 98.5% |
|  | ✓ | ✓ | 0.05 | 4452 | 225 | 24903 | 5958 | 42.8% | 99.1% | 95.2% | 80.7% |

### Compensatory mutations can identify rifampicin resistance by association with high specificity

Even when a low FRS threshold is set (e.g. 0.05) we still observe false negatives in our genotypic resistance prediction and this is unlikely to improve much upon further lowering of the FRS threshold (Fig. 2C). Another cause for false negative calls is low coverage or sequencing errors in genomic regions associated with resistance, hence it would be useful if there was redundancy when predicting resistance using genetic features. We have previously identified 51 CMs ^22^, which are, by definition, only present in resistant samples. Hence by allowing samples to be predicted as rifampicin-resistant if they contain either a RAV and/or a CM one might expect to reduce the false negative rate. To test this, we appended these 51 CMs to the WHO catalogue of RAVs for rifampicin.

Including CMs reduced the number of false negative calls from 591 to 568 when applying an FRS threshold of 0.90 to call variants (Table 1), corresponding to a 3.9% decrease. The resulting improvement in sensitivity is not significant, probably due to the small number of affected samples compared to the size of the dataset. If we repeat the experiment setting the FRS threshold to 0.05, the number of samples falsely predicted to be resistant increases from 529 to 663. Most (99%) of these erroneous calls are due to samples which, despite containing the putative CM *rpoC* F452L, are not actually resistant. If we exclude this CM *post hoc*, the specificity does not decrease when allowing CMs to predict resistance at low FRS (Table 1).

To evaluate how well CMs identify rifampicin resistance on their own, we checked the specificity of using only CMs to predict the resistance phenotype. As expected, only moderate sensitivity is achieved (42.8%, Table 1), since only a fraction of resistant samples exhibit compensation. Interestingly, the specificity of predicting resistance is higher (99.1%) than when simply applying the WHOv2 catalogue of mutations (Table 1). Overall this suggests that permitting CMs to trigger a resistance call could be useful in samples where the RAV has not been resolved, for example if the coverage or read depth is poor, but that we should not rely on them exclusively.

The combined effect of lowering the FRS threshold to 0.05 and permitting CMs to predict resistance is a reduction of false negative calls by 38.6%. Hence more than a third of the resistant samples incorrectly classified as susceptible are hereby explained and corrected (Table 1), a handy improvement.

### Heterogeneous resistant samples are less likely to show compensation

The proportion of heterogeneous samples with a compensatory mutation is significantly lower than the fraction of homogeneous samples which are compensated (43.5% v 34.2%, Fishers exact test, p-value = 3.63e-4, Fig. 3A). Since the fraction of reads supporting the CM and RAV are correlated in the compensated heterogeneous samples (Fig. 3B, p = 1.69e-22) it is likely that they are composed of a susceptible strain and a rifampicin resistant strain that has acquired a compensatory mutation.

**Figure 3:**
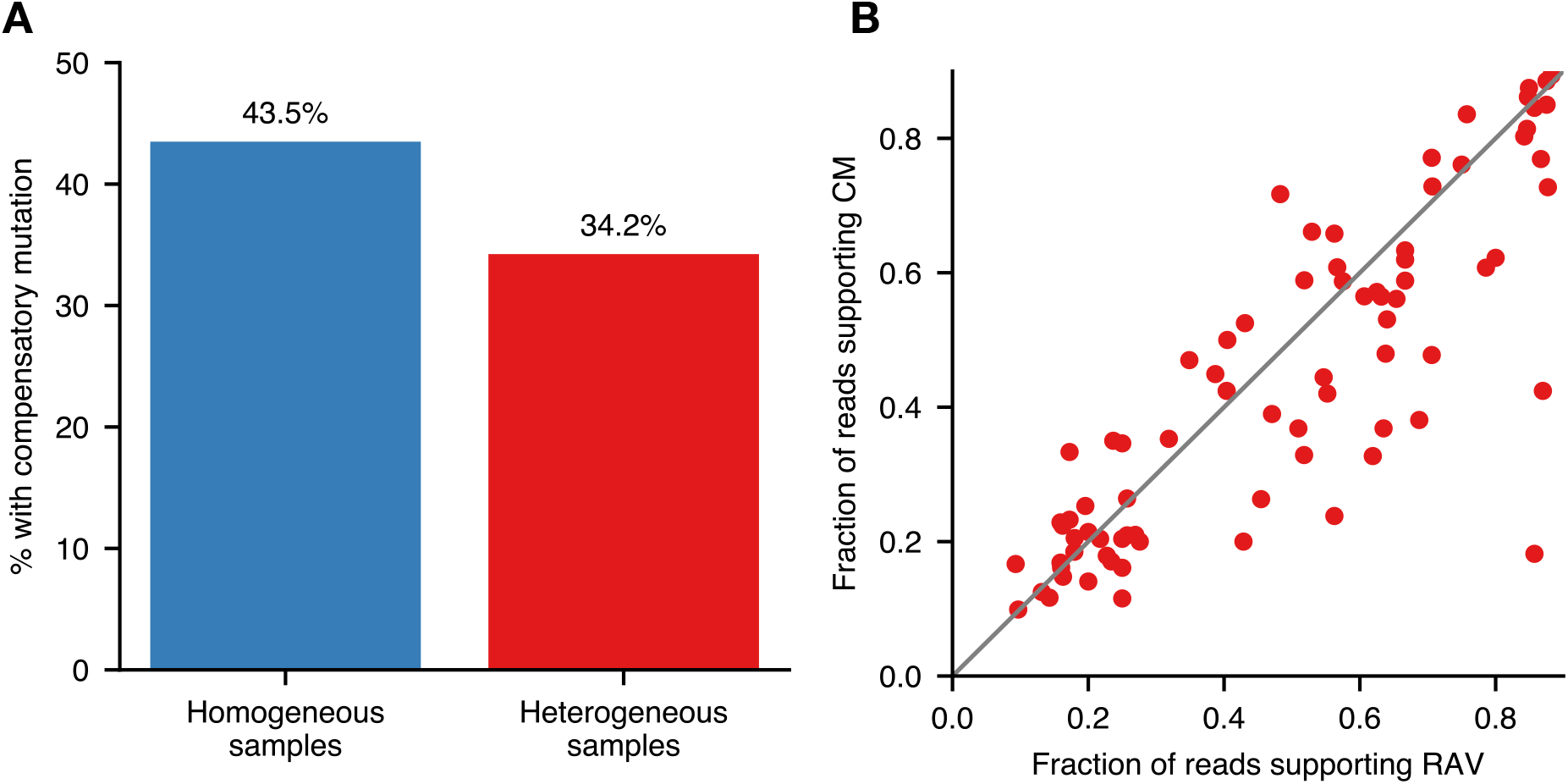
Heterogeneous resistant samples are less likely to also have a compensatory mutation (CM) than homogeneous resistant samples. **(A)** Percentage of samples showing CMs at any FRS in homogeneous vs heterogeneous resistant samples (Fishers exact p-value = 3.63e-4). **(B)** For the heterogeneous samples that are compensated, this plot shows the significant correlation between the FRS of the respective resistance and compensatory mutation (p = 1.69e-22). The grey line indicates the hypothetical line of perfect correlation.

A smaller proportion of resistant heterogeneous samples contain a *rpoB* S450L mutation; 60.5% com-pared to 68.6% in homogeneous samples. Since most compensatory mutations (96%) co-occur with this mutation, this effect would lead to the proportion of hetergeoneous samples with a CM being reduced by

4.9 percentage points to 38.6%. This is 4.4 percentage points higher than our observed fraction (34.2%) and therefore we can explain around half the difference. One possible explanation for the remainder is resistant subpopulations evolving, on average, resistance more recently than homogeneous resistant samples.

### At least 28% of heterogeneous resistant samples are the result of secondary infections

We can gain some insight into the source of resistance by examining the genetic diversity in the heterogeneous samples. If the resistant subpopulation arose by within-host evolution (Fig. 1A), one would expect to see relatively few, if any, other minor mutations in the sample, given the low mutation rate (0.04-2.2 SNPs per genome per year ^30^) of *M. tuberculosis*. A similar logic applies when a sample is taken midway through reversion of resistance following cessation of treatment (Fig. 1B); again relatively few other minor mutations would be expected. If the patient has been infected more than once (Fig. 1C), the sample will probably contain different strains and therefore we would expect a much greater number of minor mutations. This will not necessarily always be true: if the co-infecting strain is part of the same outbreak, the number of minor mutations will again be small. Only when the number of minor mutations is high, e.g. if the sample contains multiple (sub)lineages, can we draw a definite conclusion since in this case secondary infection is the only viable scenario.

To investigate this, we measured the number of minor mutations in all 248 heterogeneous resistant samples with a singular minor RAV. The latter condition is required to ensure that only a single resistant subpopulation is present. The resulting distribution is very broad; some samples have no or relatively few minor mutations, whilst others have several thousand (Fig. 4A).

**Figure 4:**
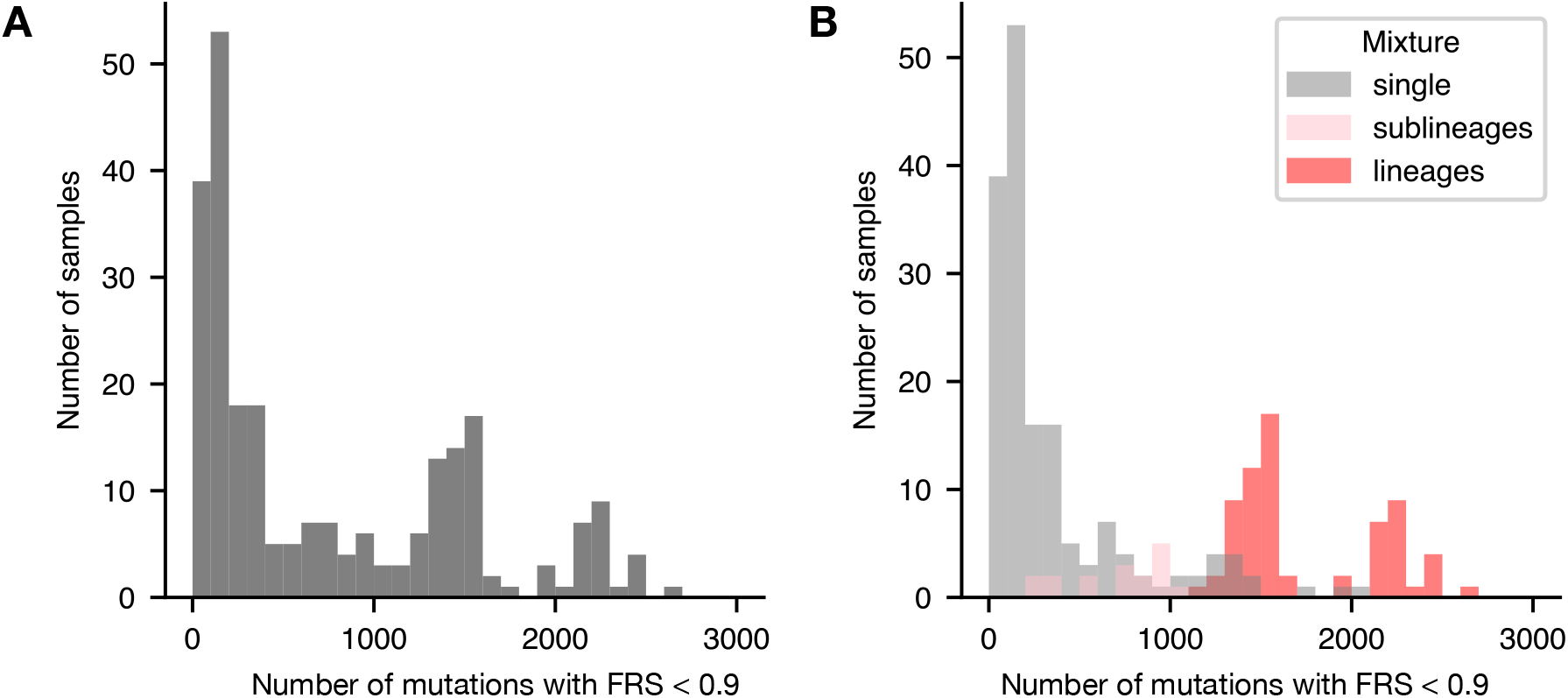
Quantifying genetic diversity within heterogeneous resistant samples using the amount of minor mutations detected. **(A)** Genetic diversity within the heterogeneous samples with resistant subpopulations, as measured by the amount of minor mutations per sample. Minor mutations are all variants in these samples which display an FRS below 0.90. **(B)** Same as in A, but the data is divided into three different batches based on the sample mixture type assigned by mykrobe (v0.12.1): A single lineage, multiple sublineages or multiple lineages per sample. All 85 samples plotted in Fig. S4.

Mapping the mixture type identified by mykrobe ^31^ (single, multiple sublineages, or multiple lineages per sample) onto the distribution neatly segregates the data into three partially overlapping subgroups (Fig. 4B). The first is the 163 samples which, according to mykrobe, consist of one and only one lineage (labelled ‘single’ in Fig. 4B). The remaining 85 samples (34 %) are assessed as either containing multiple sublineages belonging to the same lineage, or different lineages. These are labelled ‘sublineages’ and ‘lineages’, respectively, in Fig. 4B. The only plausible explanation for the latter two cases is if the patient picked up a secondary infection. To estimate how many of these were resistant at the time of infection we plotted the FRS distribution for all minor mutations present in each of the 85 samples (Fig. S4); many samples showed clear bimodality (Fig. 5A) whilst others did not (Fig. 5B). The latter case is due, we infer, to these samples containing equal parts of all lineages. For 70 of the bimodal samples the FRS of the RAV was in phase with one of the subpopulations (Fig. 5C, S4), consistent with the secondary infection being resistant at the point of transmission. The others were either clearly out of phase (Fig. 5D) or difficult to call. We conclude that at least 28% (70/248) of the heterogeneous resistant samples were the result of a secondary infection that was resistant at the point of transmission. Interestingly, there is also a significant association between a heterogeneous resistant sample having a compensatory mutation and showing more than one (sub)lineage (Fishers exact test, p-value = 2.38e-04).

**Figure 5:**
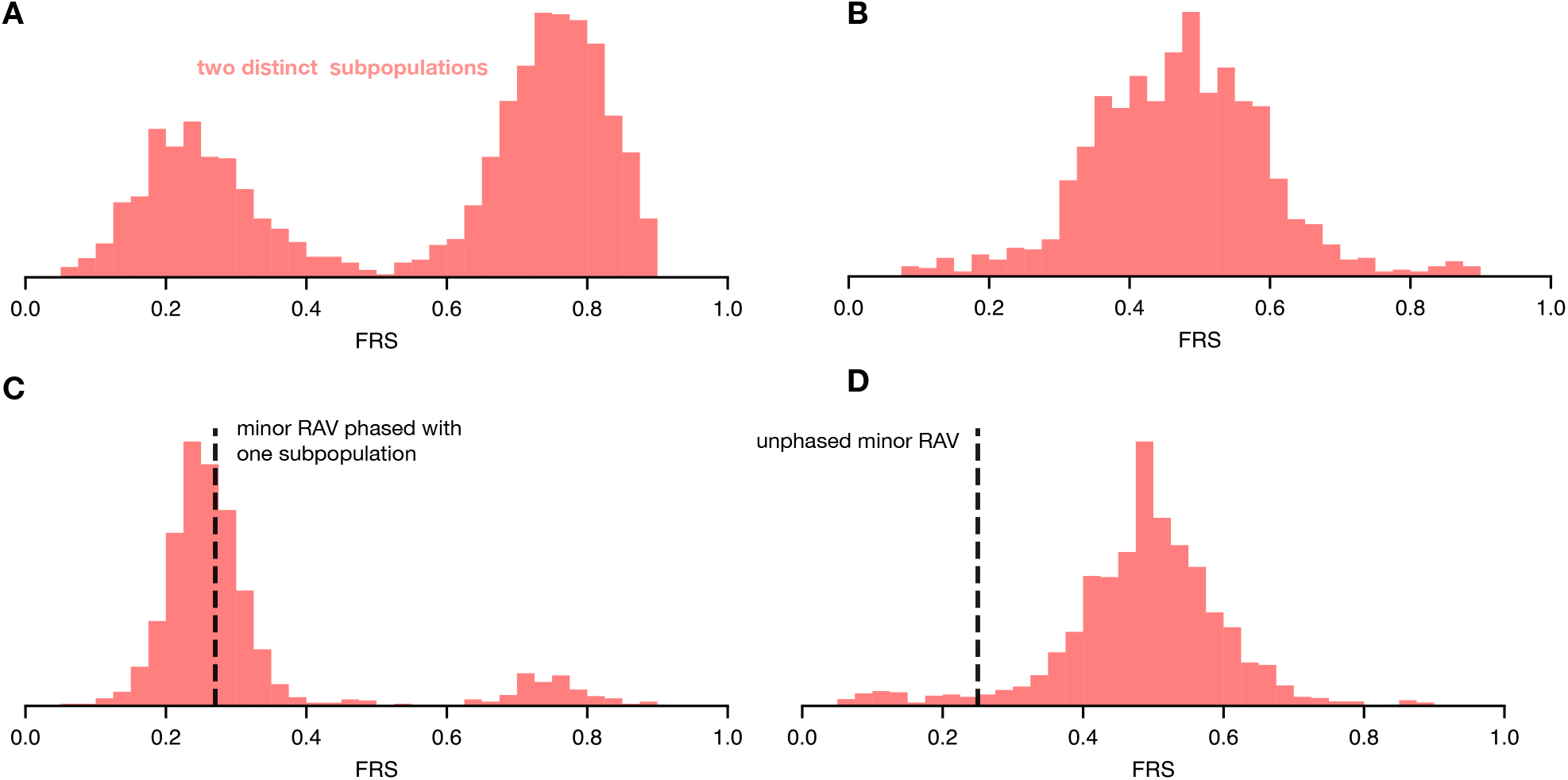
Example FRS distributions of minor mutations in heterogeneous samples with multiple (sub)lineages can show if RAVs are in phase with a subpopulation. **(A)** FRS distribution in a single sample with multiple (sub)lineages of *M. tuberculosis*. The number of (heterogeneous) mutations at each FRS are shown in histogram representation. The two subpopulations can be clearly distinguished and are around 0.2 and 0.8 in this sample. **(B)** Plot structure identical to A, but in this sample we cannot distinguish between the subpopulations due to overlapping FRS counts. **(C)** Plot structure similar to A, but additionally a black dashed line indicates the FRS of the RAV in the sample. Here it is in phase with one of the distribution maxima. **(D)** Plot structure identical to C, but the dashed line indicating the RAV is out of phase with the distribution maxima, suggesting resistance evolved after transmission.

## Discussion

We have demonstrated that it is important to consider within-host diversity when predicting drug resistance in *M. tuberculosis* using whole genome sequencing. Although heteroresistance is usually detected by targeted sequencing approaches and rapid molecular tests (e.g. Gene Xpert MTB/RIF), whole genome sequencing (WGS) approaches often fail to resolve subpopulations. But by lowering the minimum fraction of read support (FRS) required to call a resistance-associated variant (RAV) from 0.90 to 0.05, we can significantly improve the sensitivity of WGS-based genotypic resistance classification by +2.1% to 96.4%, with no detectable effect on specificity. A higher sensitivity improves any treatment decisions and leads to more effective surveillance. Importantly, it also takes rifampicin prediction on this dataset above the required minimum threshold of 95% in both sensitivity and specificity to pass the ISO standard for antimicrobial susceptibility test devices. ^10^ It would therefore seem sensible for genetic workflows that process *M. tuberculosis* samples to identify and call rifampicin RAVs if their presence is supported by just a few reads, regardless of the fraction of read support. Detecting resistant subpopulations has a greater effect in many second-line drugs. For example, it has been shown that classifying a sample as resistant to moxifloxacin based on a resistant subpopulation improves sensitivity from 85.4 % to 94.0 %. ^13^

Allowing the presence of high-confidence compensatory mutations (CMs) in the RNA polymerase to identify rifampicin resistance by association would also seem a trivial way to boost sensitivity. The improvement in sensitivity on this dataset was not significant, probably due to the moderate read depth (Fig. S3) ensuring there were relatively few resistant samples with a compensatory mutation where the resistance mutations could not be resolved but the CMs could. In scenarios with much lower read depth (for example in multiplexed samples) allowing compensatory mutations to predict rifampicin resistance could provide a necessary boost to the sensitivity. In future work we will investigate the read depths at which it becomes beneficial to allow CMs in addition to RAVs to call rifampicin resistance by randomly down-sampling sequencing reads.

As noted above, it is vital that any list of compensatory mutations is accurate if they are to be used to infer resistance. ^22^ The mutation *rpoC* F452L appears to not be resistance-associated when present at lower FRS, which could either be an error or point to an inverse causal relationship between this CM and the presence of RAVs. This raises the question of whether F452L should be considered a CM or rather provides an advantageous genetic background for the acquisition of rifampicin resistance, by raising fitness *prior* to resistance emergence. Whilst speculation, this is supported by the observation that *rpoC* F452L was shown to stabilise the open promoter complex and increase the elongation rate of the *M. tuberculosis* RNA polymerase not only in the resistant *rpoB* S450L mutant, but also in the wild type. ^32^

Overall, reducing the minimum FRS threshold to 0.05 and permitting the presence of CMs to predict rifampicin resistance raises sensitivity to 96.5%. The remaining discordance can likely be explained by several factors: there may be additional, unknown rare resistance mutations in the RNA polymerase that have not been classified as such by the second edition of the WHO resistance catalogue. Tackling this problem is non-trivial but could involve the use of machine learning models trained to predict the effect of individual *rpoB* mutations. ^33^ There are almost certainly errors in the laboratory processes, such as mislabelling or measurement error, that we can minimise but not eradicate. More straightforwardly, phenotypic methods have been shown to detect 1% heteroresistance ^14^ which is more sensitive than permitted by the mean read depths of this dataset (Fig. S3): lowering our threshold from 0.05 to 0.01 would require sequencing to greater depth. Even smaller subpopulations may be relevant; it has been shown that genetics can detect microheteroresistance (0.001 *<* FRS *<* 0.05), and this could be indicative of the subsequent development of phenotypic drug resistance in clinical practice. ^34^

We can also gain insight into the timing and origin of resistance to rifampicin by examining the samples with resistant subpopulations. Around half (53%) of the significant reduction in the proportion of compensated heterogeneous resistant samples can be explained by the lower prevalence of the *rpoB* S450L mutation. Explanations for the remaining fraction include resistant heterogeneous samples having acquired resistance more recently and the varying fitness costs of different RAVs. The number of other putative minor mutations in the heterogeneous samples hints at the probable sources of resistance. Our data are consistent with secondary infection, as opposed to within-host evolution, being the root cause of rifampicin resistance in at least 28% of the heterogeneous samples. This is consistent with a longitudinal study which found that out of 107 patients with recurrent tuberculosis, 54 (50.5%) had become resistant to rifampicin via a secondary infection ^35^. The striking level of genetic diversity detected within our samples is only achievable if the patient was subject to a secondary infection, which likely introduced the resistant subpopulation. The significant association of CMs with samples containing multiple (sub)lineages indicates these samples are also more likely to be compensated. This is consistent with the secondary infection scenario, since the infecting strain could already have been compensated before being transmitted.

Stepping back, it is not surprising that the CRyPTIC dataset contains genetically heterogeneous *M. tuberculosis* samples due to how clinical samples are grown and colonies extracted. For example, aliquots taken from positive MGIT tubes will contain multiple ‘crumbs’ and therefore multiple colonies. This is in sharp contrast to laboratory practices for other pathogens where single colony picks from solid media are commonplace. One should therefore, perhaps, consider Mycobacterial genetic sequencing as inherently metagenomic.

In conclusion, this study has shown how to improve the sensitivity of WGS-based rifampicin resistance prediction by including subpopulations, compensatory mutations, and population diversity. It also provides useful information on how resistance emerges clinically. In general, these findings will facilitate improved diagnostic strategies and more effective management of drug-resistant tuberculosis.

## Acknowledgements

We would like to thank Professors Tim Peto, Nicole Stoesser and David Eyre for helpful suggestions.

## Transparency declaration

PWF receives consultancy fees from the Ellison Institute of Technology, Oxford Ltd.

