## Supplemental Information for "Subpopulations in clinical samples of *M. tuberculosis* can give rise to rifampicin resistance and shed light on how resistance is acquired"

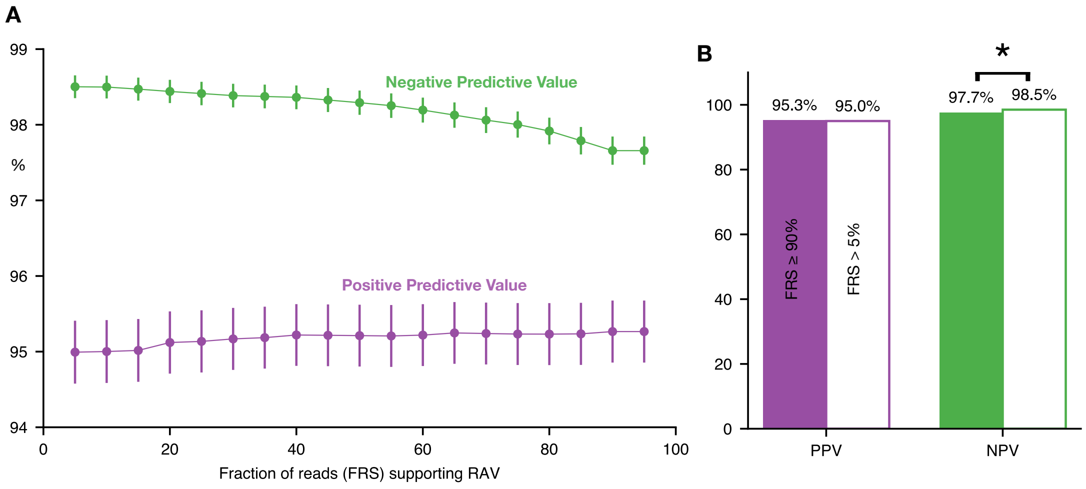


**Figure S1: Classifying samples containing a subpopulation with a resistance-associated variant (RAV) as resistant improves the NPV with no effect on PPV. (A)** Decreasing the FRS threshold required to support a variant call that is a known RAV increases NPV with little effect on PPV. The slopes of a linear regression for NPV and PPV are -0.009 and 0.003, respectively. Error bars (95% confidence limits) are plotted as calculated via the binomial proportion. **(B)** The NPV is significantly improved if the FRS threshold is lowered from 90% to 5% (z-test, p-value 5e-12). There is no significant difference in the change in PPV. For clarity error bars are not plotted.

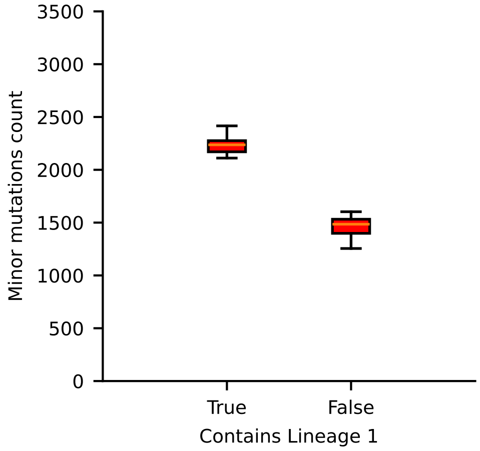


**Figure S2**: **Samples that show multiple lineages and contain at least one ancient *M. tuberculosis* lineage (Lineages 1, 5-7) have more minor mutations (mutations with $<$ 0.90 FRS) than those samples with exclusively modern lineages (Lineages 2-4).** The average minor mutation count for samples with Lineage 1 is 2598, while for exclusively modern lineage samples it is 2240.

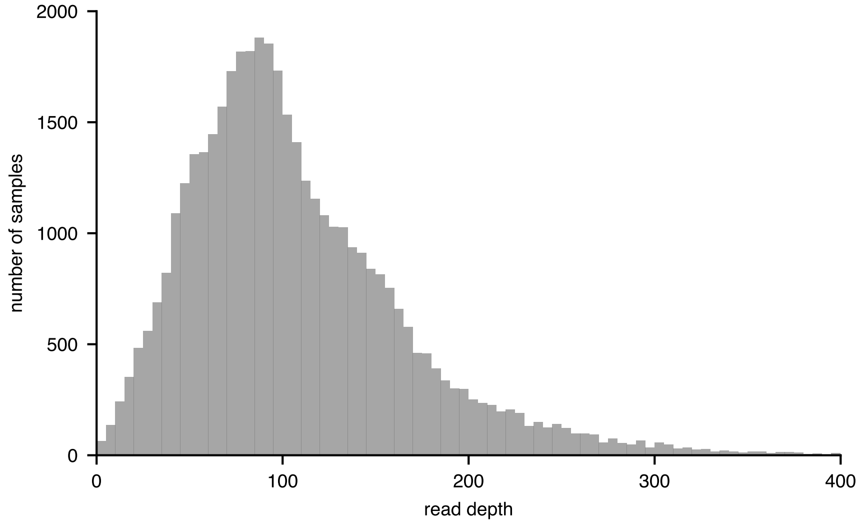


**Figure S3**: **The samples have a range of mean read depths**. For the 35,538 samples with both WGS and pDST data the mean and modal depth are 108 and 96, respectively. A majority of samples (79.8%) have a mean read depth of at least 60 which permits a minor variant at an FRS of 0.05 to be detected (as three reads are required). The chance of detecting a minor variant at an FRS of 0.01 in this dataset is low because only 1.2% of samples have a mean read depth of 300 or greater.

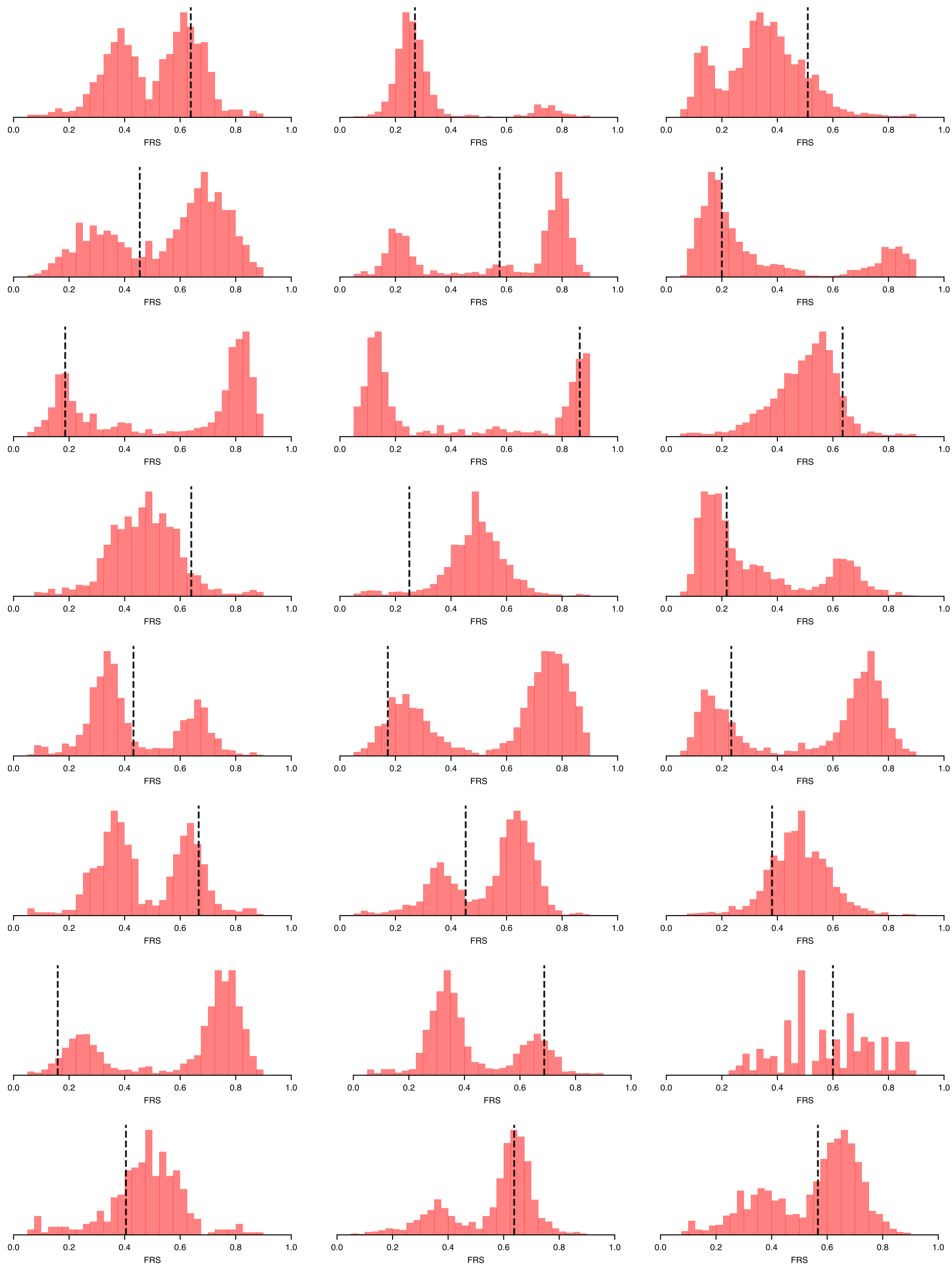


**Figure S4**: (related to Figure 5) Distributions of the fraction of reads (FRS) supporting all detected minor mutations in the 85 heterogeneous resistant samples that do not have a single lineage (Figure 4). Samples typically show unimodal, bimodal or complex behaviour. The FRS of the detected resistance-associated variant is annotated via a dash black line. If it coincides with one of the subpopulations it is in phase and can be reasonably assumed to not have evolved since transmission. Samples 1-24 (out of 85) shown here.

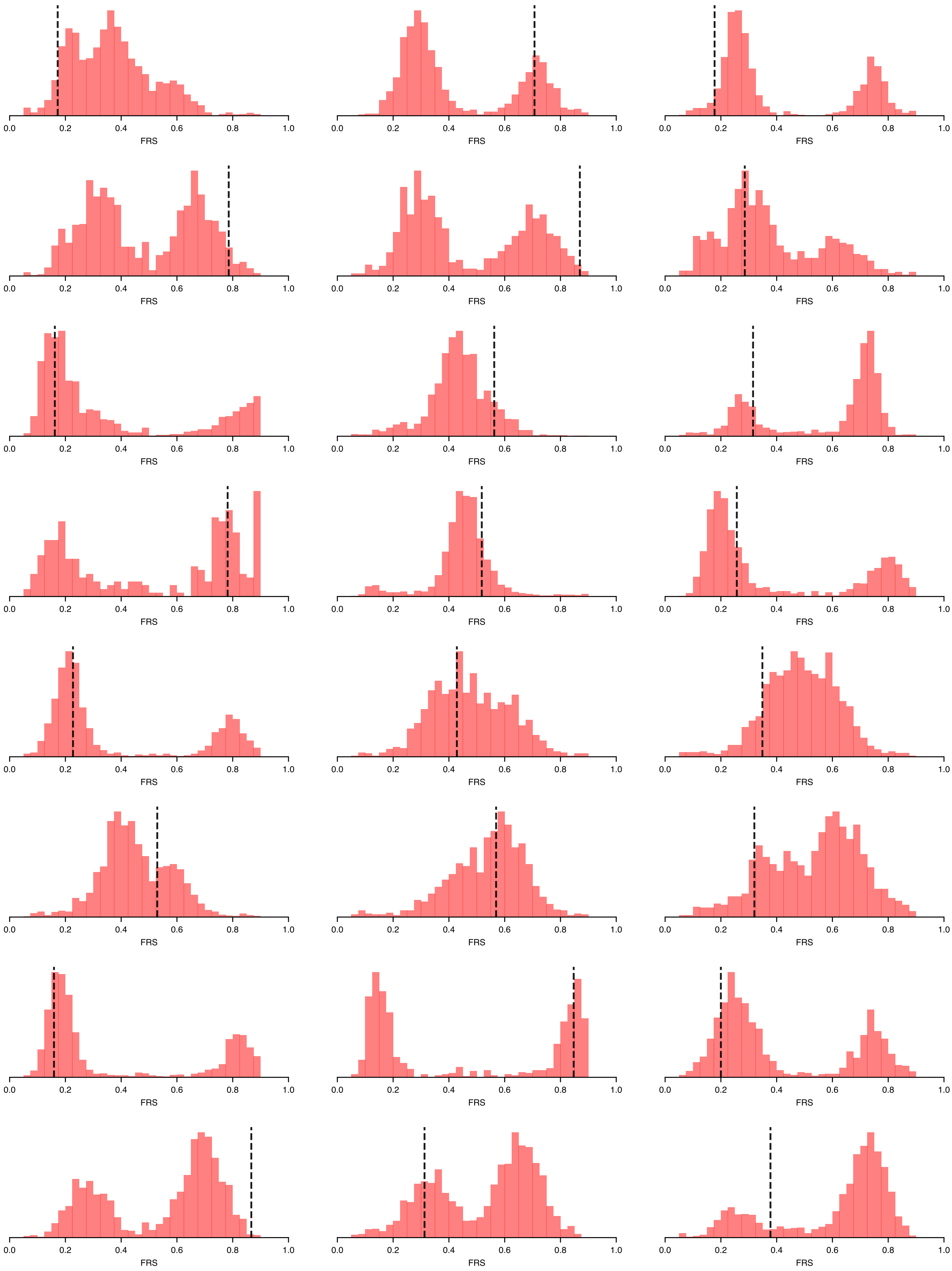


**Figure S4 continued**: Samples 25-48 (out of 85) shown here.

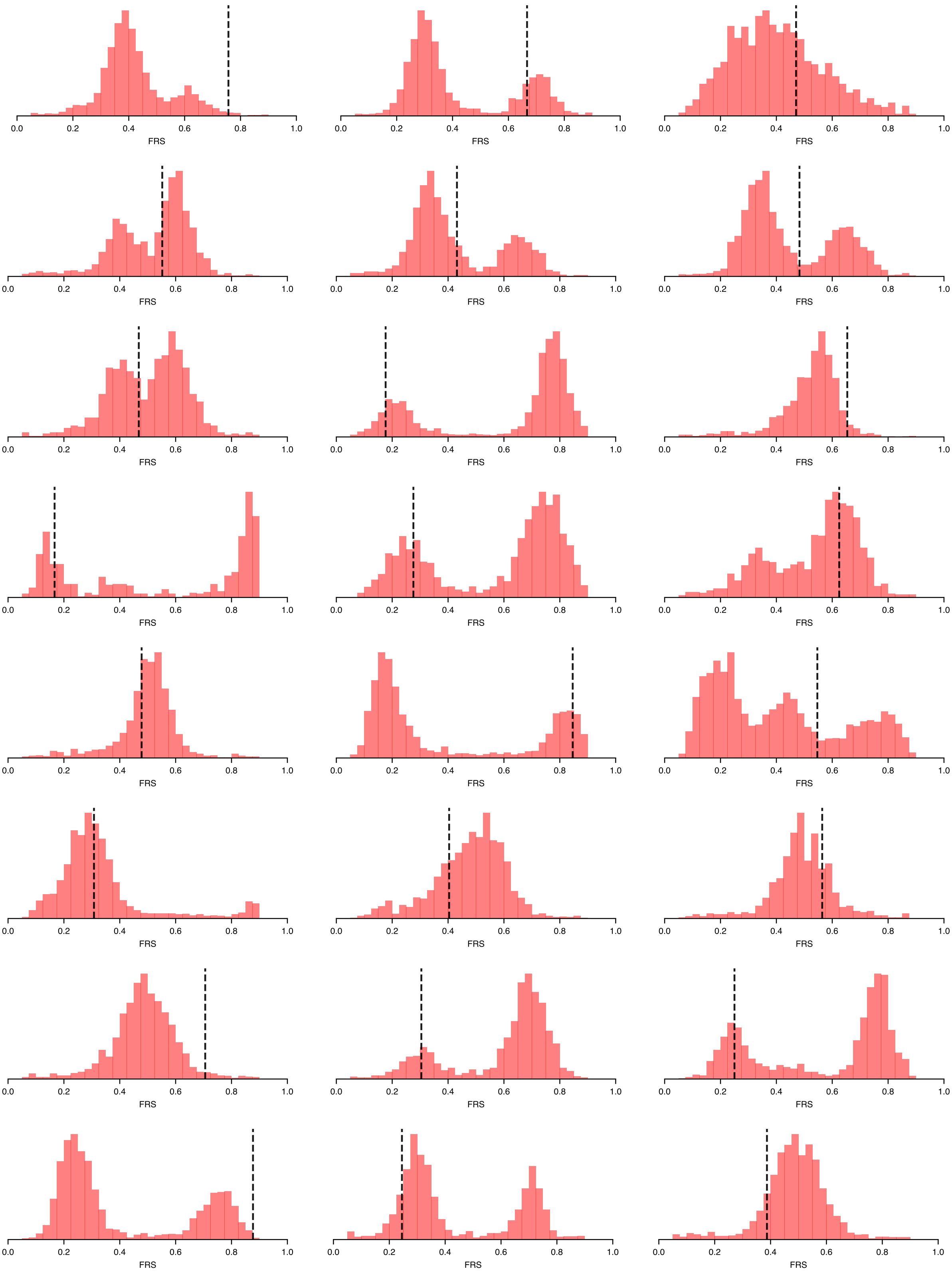


**Figure S4 continued**: Samples 49-72 (out of 85) shown here.

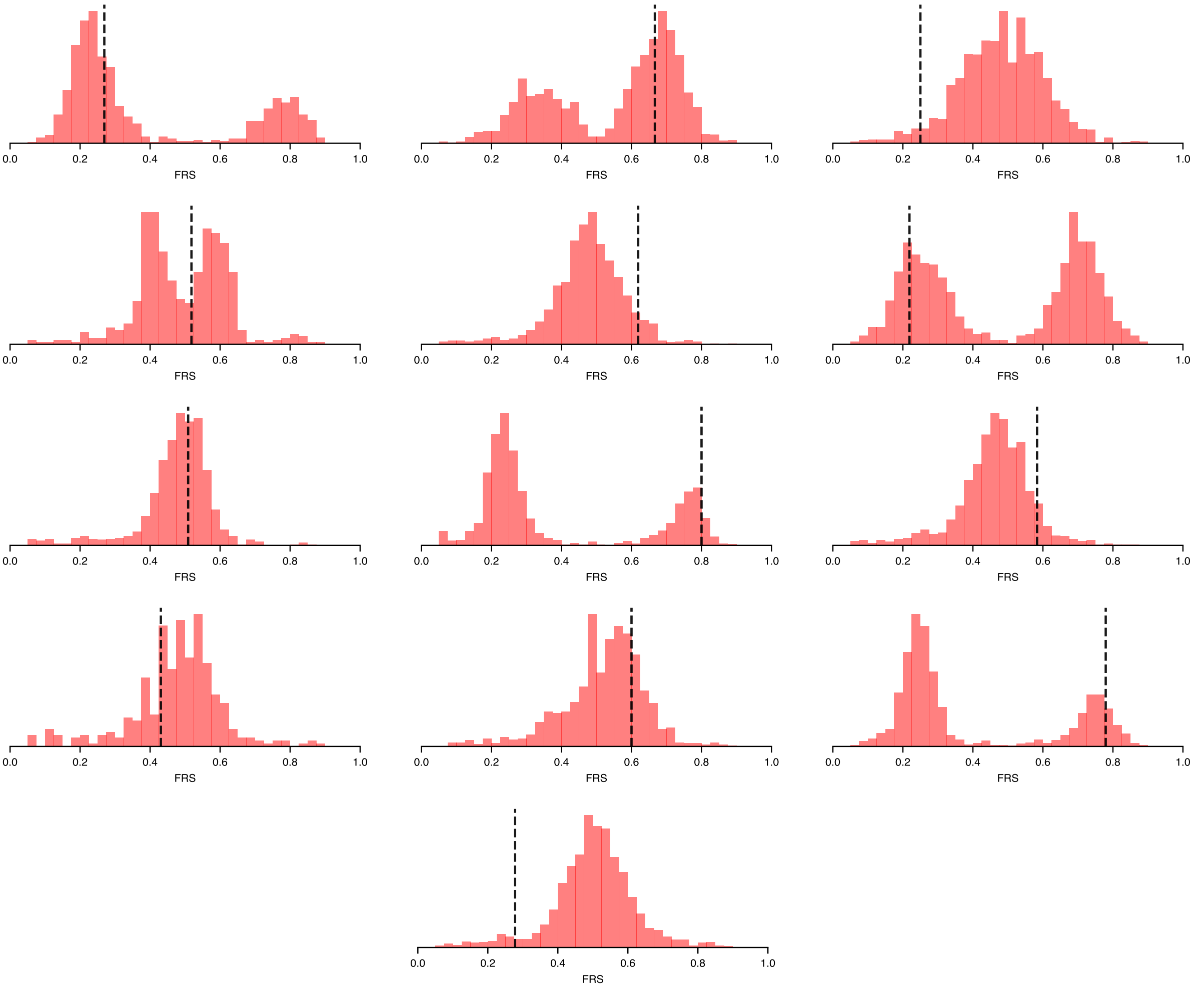


**Figure S4 continued**: Samples 73-85 (out of 85) shown here.
